# Large-Scale plurimodal networks common to listening, production and reading word-lists: an fmri study combining tasks-induced activation and intrinsic connectivity in 144 right-handers

**DOI:** 10.1101/382960

**Authors:** I Hesling, L Labache, M Joliot, N Tzourio-Mazoyer

## Abstract

Even if speech perception and production have been revealed to share a common recruitment of both discrete auditory and motor areas, this overlap being also common to reading and listening, no study has investigated the involvement of larger networks in the three tasks yet. So, we first identified the multimodal bilateral brain areas conjointly activated and asymmetrical during listening, production and reading of word-list using fMRI in 144 healthy right-handers (27 years ± 6 years). Such a selection made it possible to unravel 14 regions of the left hemisphere including motor, premotor and inferior parietal cortical areas. On the right, 7 regions were selected, including the posterior Human Voice Area (pHVA). To characterize the network organization within these 21 regions, we then analysed resting-state functional connectivity in 138 of the same participants. It allowed us to segregate a network of executive areas in relation with task completion from a bilateral WORD_CORE network composed of (1) all left areas supporting the action-perception cycle, in which articulatory gestures are the central motor units on which word perception, production, but also reading, would develop and act together according to the motor theory of speech; (2) the right pHVA, acting as a prosodic integrative area, underpinning the intertwining across hemispheres between prosodic (pHVA) and phonemic (left SMG) processing. The present results show that word processing, whatever the language modality involved, is based on a network of plurimodal areas hosting processes specific to each hemisphere and on their cooperation built upon synchronisation at rest.

## Introduction

Language is one of the most important and specific cognitive abilities of human beings. According to Saussure (Saussure, 1975), language is a universal structure encompassing the abstract, systematic rules and conventions of a unifying system, which is independent of individual users, whereas speech is the personal use of language, thus presenting many different variations such as style, grammar, syntax, intonation, rhythm, pronunciation. Even if neuroimaging studies of language have demonstrated a bilateral involvement, language is implemented in large areas located along the left sylvian fissure (Vigneau et al, 2006, Vigneau et al, 2011). More specifically, word processing is underpinned by core language areas located in auditory, visual and motor cortical areas of the left hemisphere depending on the type of language activity (Price 2012, 2010). Concerning the left hemisphere setup of language areas, there exist two divergent theories about the relation of speech perception and production to language. The first one, coined the horizontal view, proposes that the elements of speech are sounds that rely on two separate processes (one for speech perception, the other one for speech production) which are not specialized for language until a cognitive process connects them to each other and then to language (Fodor, 1983). The second theory, coined the vertical view (or motor theory of speech perception), posits that speech elements are articulatory gestures serving both speech perception and production processes which are immediately linguistic, thus requiring no cognitive process (Liberman and Whalen, 2000). At the cerebral level, in line with the motor theory of speech perception, the existence of a bilateral dorsal-ventral model of speech processing, with a preferential leftward involvement, has been widely admitted (Binder et al., 1996; Hickok and Poeppel, 2004; Rauschecker and Tian, 2000). This model has posited: (i) a dorsal pathway, i.e., the “where stream,” where an acoustic-phonetic-articulatory transformation linking auditory representations to motor representations is reported to occur in superior temporal/parietal areas and ultimately in frontal areas (Buchsbaum et al., 2001); and (ii) a ventral pathway, i.e., the “what stream”, where speech-derived representations interface with lexical involve the superior, middle, and inferior temporal gyri (Binder et al., 2000; Hickok and Poeppel, 2000). Interestingly, concerning the dorsal pathway, the postulate of the existence of an auditory-motor system (Hickok and Poeppel, 2000), has been supported by studies aiming at examining the role of motor areas in speech perception. Hence, an fMRI study has revealed that listening to syllables and producing the same syllables lead to a common bilateral network encompassing a superior part of the ventral premotor cortex, suggesting the existence of a common phonetic code between speech perception and production (Wilson et al., 2004). Furthermore, another study has not only put forward that the cortical motor system is organized in a somatotopic way along the precentral cortex, the lip area being superior to the tongue area, but has also revealed that these precentral regions are consistently activated by syllable articulation and syllable perception, hence demonstrating a shared speech-sound-specific neural substrate of these sensory and motor processes (Pulvermüller et al., 2006). These findings have been supported by a meta-analysis revealing that in right-handers, activations of the posterior part of the frontal lobe distributed along the precentral gyrus are strongly left lateralized during both production and auditory tasks at the word or syllable level, together with the involvement of the SupraMarginal Gyrus (SMG) (Vigneau et al., 2006). Moreover, a recent MEG study has reported a synchronisation between anterior motor regions involved in syllables articulation and posterior regions involved in their auditory perception during both their production and perception (Assanéo & Poeppel, 2018).

Though mastered more belatedly, human beings have developed ways of using language through other sensory modalities, such as the visual system in the case of reading. Accurate perception and production of speech sounds are essential for learning the relationship between sounds and letters. Phonological awareness, i.e., the ability to detect and manipulate speech sounds, or phonemes, is the best predictor of reading ability. Reading is based on both the ability to hear and segment words into phonemes and then to associate these phonemes with graphemes, the mapping of orthographic to phonological representations during reading being intrinsically cross-modal (Mc Norgan et al, 2014). Research has revealed that a phonological processing deficit underlies reading difficulties in dyslexic children, establishing a link between perception and reading abilities (Gillon 2004). In the case of disorders of oral language development, Specific Language Impairment, SLI, is the most frequently studied developmental disorder. Children with SLI have been reported to present impairments in phonological processing, be it phonological awareness or phonological memory, establishing a link between production and reading abilities, whose neural support is still under question (Catts et al, 2005). Different studies examining word processing cerebral networks common to the auditory and visual modalities have revealed the supramodal implication of anterior regions (supplementary motor area - SMA -, prefrontal, premotor and inferior frontal gyrus) whereas variations have been observed in the temporal lobe according to the language task (Booth et al., 2002a, 2002b; Buckner et al., 2000; Chee et al., 1999), making it difficult to conclude on the existence of a common antero-posterior network for plurimodal word processing. Note that concerning semantic processing, one study, dealing with production and reading in four languages revealed a common bilateral network involved by both components (Rueckl et al, 2015).

Importantly, even if less investigated, the first phase of speech acquisition in newborn babies is perceptual as the infant hears others’ vocalizations, highlighting the importance of prosodic areas in speech processing. Speech prosody, i.e., the musical aspects of speech, is an early-developing component of speech, which could be compared as a musical stave, upon which phonemes would be placed (Locke, 1990). This perceptual phase is crucial considering the inability to learn spoken language or even babble normally when infants are born deaf (Oller and MacNeilage, 1983), or in the case of wild children (Curtiss, 1977). Lesional studies have revealed that tonal prosodic brain areas are located in the right hemisphere along the STS, including the posterior Human Voice Area (pHVA,) underlying the potential role of these right hemispheric regions during development. The second phase of speech acquisition is production. In fact, children would master the prosodic dimension, the expressivity dimension, before producing their first words, i.e., the expression dimension (Bever et al, 1971). Production develops through the process of imitation, highlighting that prosodic processing is one element of the construction of a strong dependency between perception and production through development. This is illustrated, for example, by persisting difficulties in speech production encountered by infants who were tracheotomized at the time at which they should have normally babbled (Locke and Pearson, 1990). Interestingly, meter in speech, whose acoustic correlate is stress, has been revealed to be important for both speech perception and production (Jusczyk et al, 1993). Once the metrical rules, which provide important cues for speech segmentation within the continuous speech stream, have been acquired, speech meter contributes to phonological (Pitt & Samuel, 1990), semantic (Schwartze et al, 2011) and syntactic (Roncaglia-Denissen et al, 2013) processing. Musical rhythmic priming, using meter, has been revealed to enhance phonological production in hearing impaired children, due to an enhanced perception of sentences (Cason et al, 2015). Furthermore, in the context of speech rehabilitation therapies, musical rhythm has been revealed to be a fluency-enhancing tool (Thaut, 2013). More generally, the prosodic dimension of speech is used to restore the speech of Broca’s aphasic patients and the term Melodic Intonation Therapy was coined to refer to this technique based on the use of melody and singing which would be core musical elements engaging predominantly the right hemisphere (Thaut & McIntosh, 2014).

Taken together, and considering the specialization of the left and right hemispheres for different aspects of language and speech processing, these results suggest the existence of plurimodal networks common to the production, listening and reading of word-lists. For example, studies on split-brain patients have demonstrated a strict leftward lateralization concerning phonological processing, split-brain patients’ right hemisphere lacking categorical perception of phonemes (Gazzaniga 2000, Sidtis et al., 1981). Such a leftward lateralization was confirmed by studies using the Wada test procedure (Dym et al., 2011) and the leftward asymmetry of the audio-motor loop measured with functional imaging actually supports the left hemisphere specialization for the phonological processing of speech dimension (Vigneau et al. 2005 and 2008). Another example is the right STS specialization for tonal processing, also evidenced by neuroimaging as a rightward asymmetry of activation during such sounds processing (Zatorre & Belin, 2001). Considering the importance of prosody in language development, we hypothesized that in addition to the left hemisphere participation, right hemispheric regions hosting the tonal dimension of speech prosody would be involved in the 3 tasks, i.e., production, perception and reading tasks (Beaucousin et al., 2007; Belin et al., 2004; Sammler et al., 2015). Moreover, since the complete development of speech in literates leads to the mastering of written speech, we expected that the core word areas that developed conjointly in the 3 modalities would include some visual areas as being part of a large-scale plurimodal network underpinning word processing.

To achieve the identification of plurimodal large-scale networks for word-list processing, we selected 144 right-handers from the BIL&GIN database (Mazoyer et al., 2016) because we have shown that almost all right-handers have a left hemisphere dominance for language (Tzourio-Mazoyer et al., 2016, Zago et al., 2017) to: (1) identify brain regions conjointly activated and asymmetrical during word production, word perception and word reading in each hemisphere; (2) conduct a comprehensive investigation of how these areas are modulated according to the task to elaborate their function/role; (3) identify the networks at play within the areas commonly shared by the three tasks based on the calculation of the intrinsic connectivity, in the same participants.

## Material and methods

### Participants

The present study included a sample of 144 right-handers balanced for sex (72 women) from the BIL&GIN database, which is a multimodal imaging/psychometric/genetic database specifically designed for studying the structural and functional neural correlates of brain lateralization (Mazoyer et al., 2016). Participants were selected as having French as their mother tongue and were free from developmental disorders, neurological and psychiatric history. A local ethic committee (CCPRB Basse-Normandie, France) approved the experimental protocol. Participants gave their informed, written consent, and received an allowance for their participation. All subjects were free of brain abnormalities as assessed by an inspection of their structural T1-MRI scans by a trained radiologist.

The mean (± standard deviation) age of the sample was 27 years ± 6 years (range: 19-53) and the mean level of education (corresponding to the number of schooling years sincethefirstgradeofprimaryschool)was16±2years(range: 11-20) corresponding to four years at the university level. Handedness was self-reported by the subjects and their Manual Lateralization Strength (MLS) was assessed using the Edinburgh inventory (Oldfield, 1971) whose values ranged from −100 to +100. Average MLS values were 93.48 (S.D. = 11.49) for the subjects included in the present study.

### Functional imaging

#### Paradigm of the word-list tasks

Language mapping was assessed in the three language tasks at the word level during which participants had to covertly generate (PROD), listen (LISN) or read (READ) lists of words (Word-List). These tasks were part of a run that alternated these word tasks with sentence tasks (see Labache et al, 2019 for the complete methodology).

The stimuli were lists of over-learnt words, such as months of the year, making it possible to decrease the weight of lexico-semantic and syntactic processing and enhancing phonetic encoding, phonology, articulation and word retrieval while inducing a prosodic processing due to the specific metrics of lists. For each task, the participants were shown a scrambled drawing during 1 s, immediately followed by a central fixation crosshair (see Figure 1). While fixating the cross, the subjects performed either the listening, production or reading Word-List task and had to click on the response pad after task completion. Then a low-level reference task followed each event (sentence or Word-List), consisting in sustaining visual fixation on the central cross and pressing the pad when the fixation cross was switched to a square (both stimuli covering a 0.8°×0.8° visual area). This second part of the trial, that lasted at least half the total trial duration, aimed at refocusing the participant attention on a non-verbal stimulus and to control for the manual motor response.

**Fig. 1.**
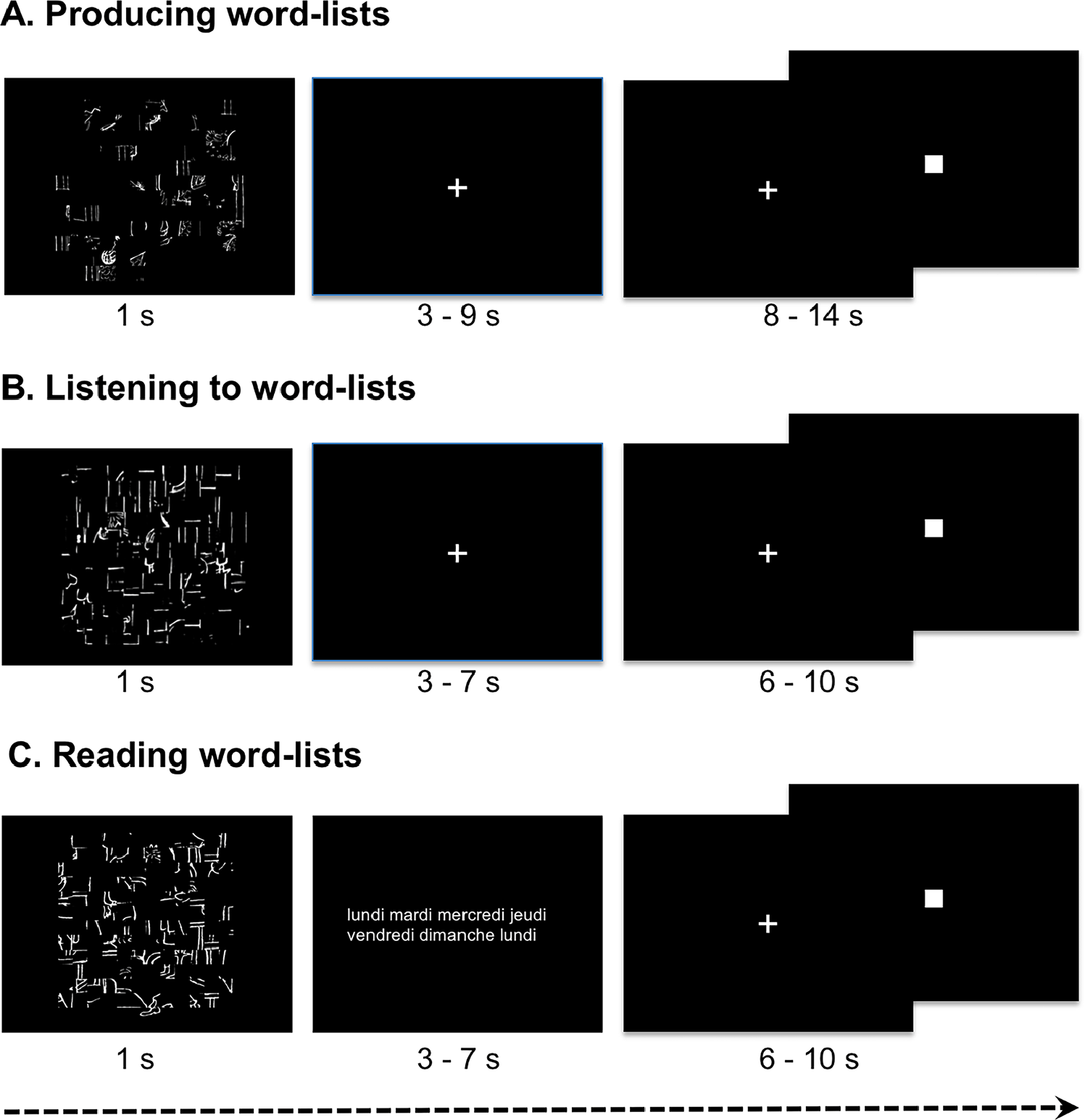
Description of the paradigm for one event of each Word-List task. In the three conditions, an event started by the presentation of a scrambled picture during one second, followed by a central cross participants were instructed to fixate while they were producing the list of months of the year (A), or listening to a list of words (B). During the reading run instead of a cross fixation, lists of words (either weeks, hours, seasons, days, months) were presented. They had to click when they had finished to produce, listen to or read and next they had to indicate by clicking when the central cross that reappeared was changed into a square. For this figure, for reading convenience, stimuli were zoomed as compared to the presentation of stimuli during the scanning session.

The design was almost identical for the 3 tasks. During PROD, the participants were asked to covertly generate the list of the months of the years from January to December during each of the ten Word-List trials lasting 18-s. During LISN, they were instructed to listen to thirteen 14-s trial of list of the months, days of the week and/or seasons. During READ, they were asked to read list of days of the weeks, months, and/or seasons flashed on the screen and consisting in thirteen trials lasting 14-s.

#### Image acquisition and pre-processing

##### Structural imaging

Structural images were acquired using the same 3T Philips Intera Achieva scanner including high-resolution T1-weighted volumes (sequence parameters: TR, 20 ms; TE, ms; flip angle, 10°; inversion time, 800 ms; turbo field echo factor, 65; sense factor, 2; field of view, 256×256×180 mm^3^; isotropic voxel size 1 × 1 × 1 mm^3^). For each participant, the line between the anterior and posterior commissures was identified on a mid-sagittal section, and the T1-MRI volume was acquired after orienting the brain in the bi-commissural coordinate system. T2*-weighted multi-slice images were also acquired [T2*-weighted fast field echo (T2*-FFE), sequence parameters: TR = 3,500 ms; TE = 35 ms; flip angle = 90°; sense factor = 2; 70 axial slices; 2 × 2 × 2 mm^3^ isotropic voxel size].

##### Task-induced image acquisition and analysis

The functional volumes were acquired as T2*-weighted echo-planar EPI images (TR = 2 s; TE = 35 ms; flip angle = 80°; 31 axial slices with a 240 × 240 mm^2^ field of view and 3.75 × 3.75 × 3.75 mm^3^ isotropic voxel size). In the three runs, 192, 194 and 194 T2*-weighted volumes were acquired for the Word-List production, listening and reading tasks, respectively.

For each participant, the pre-processing of T2*-weighted echo-planar EPI images included (1) the T2*-FFE segmentation into three brain tissue classes: grey matter, volume rigid registration to the T1-MRI; (2) the T1-MRI white matter and cerebrospinal fluid; and (3) the T1-MRI scans normalization to the BIL&GIN template including 301 volunteers from the BIL&GIN database (aligned to the MNI space) using the SPM12 “normalise” procedure with otherwise default parameters. For each of the three fMRI runs, data were then corrected for slice timing differences and to correct for subject motion during the runs, all the T2*-weighted volumes were realigned using a six-parameter rigid-body registration. The EPI-BOLD scans were then registered rigidly to the structural T2*-FFE image. The combination of all registration matrices allowed for warping the EPI-BOLD functional scans to the standard space with a single trilinear interpolation.

The analysis included first, the application of a 6-mm full width at half maximum (Gaussian filter) to each functional volume of each run. Global linear modelling (statistical parametric mapping (SPM), http://www.fil.ion.ucl.ac.uk/spm/) was then used for processing the task-related fMRI data. For each participant, BOLD variations corresponding to each Word-List task versus cross change detection belonging to the same run were computed [Word-List production (PROD), Word-List reading (READ), and Word-List listening (LISN)].

##### Homotopic regions of interest analysis

Since the brain presents a global torsion, the Yaklovian torque, which makes difficult a point-to-point correspondence between cortical areas that are functionally homotopic (Toga and Thompson, 2003), the use of flipped images to calculate asymmetries appears problematic since it cannot ensure that the flipped regions correspond to the equivalent of the other hemisphere. A new atlas AICHA, based on resting state fMRI data and composed of homotopic functional Regions Of Interest (hROIs), has been devised to circumvent this latter problem and is thus suited for investigating brain hemispheric specialization and lateralization, allowing for determining right and left hemispheric contribution in language and for computing functional asymmetries in regions having equivalent intrinsic connectivity (Joliot et al., 2015).

BOLD signal variations were thus calculated for the right and left hROI BOLD signal variations for each of 185 pairs of functionally defined hROIs of the AICHA atlas (Joliot et al. 2015) adapted to SPM12 in the 3 contrast maps (defined at the voxel level) of PROD, LISN and READ.

### Part 1. Identification and Characterization of hROIs exhibiting both activation and asymmetry in all three tasks

To complete the identification of language areas underpinning production, listening and reading tasks at the word level, we first searched for hROIs that were both significantly conjointly activated and significantly asymmetrical on average among the 144 participants during the PROD, LISN and READ tasks for each hemisphere. The idea behind the conjunction of activations and asymmetries during the 3 tasks is to be selective enough to present brain areas specifically lateralized during the tasks. In a second step, we described the variation in activity and in asymmetry in each hemisphere for the selected regions in order to obtain information on their functional nature from their modulation by the task component.

### Statistical analyses

#### hROIs selection

Using JMP14 (http://www.jmp.com, SAS Institute Inc., 2018), conjunction analysis was conducted to select the left hemispheric hROIs exhibiting BOLD signal variations that were both significantly positive and significantly larger than that of their right counterparts in all three tasks. An hROI was selected whenever it was significantly activated in each of the three task contrasts using a significance threshold set to p < 0.05 per contrast. The significance threshold for the conjunction of activation in the three tasks was thus 0.05 × 0. 05 × 0.05 = 1.25 × 10^−4^. The second criterion for hROI selection was the existence of a significant leftward asymmetry in each of the three task contrasts, the threshold of significance of this second conjunction being 1.25 × 10^−4^. Finally, to be selected, a given hROI had to fulfil both criteria, the overall significance threshold for the conjunction of conjunction analyses was 1.5 × 10^−8^ = (1.25 × 10^−4^)^2^. This procedure was conducted separately for the left and for the right hemisphere.

#### Characterization of hROIs activations and asymmetries across tasks

Taking right and left hROIs separately, we tested the existence of significant effects of Task (PROD, LISN, READ) on the selected hROIs, as well as an effect of Side (left or right), and their interactions with hROIs using two repeated-measures linear mixed-effects models. The analysis was conducted in the 144 individual entering the variable Subject as a random effect.

Two-sided Tukey’s range tests on the mean activation or asymmetry values were then completed for the left (14 hROIs) and right (7 hROIs) hROIs in order to identify the origins of significant effects and interactions found for each linear mixed-effects models.

## Results

### Tasks performances during the scanning session

The mean response time taken for the covert generation of the months of the year was 5261 ms ± 1092 ms, (range: 2836-7360). The mean response time taken for reading the months of the year, days and seasons was 4405 ms ± 603 ms, (range: 2703-5681). The mean response time taken for listening to the months of the year, days and seasons was 486 ms ± 101 ms, (range: 282-794). As the mean response time was calculated after the delivery of the stimulus, i.e., after 4386 ± 484 ms of Word-Lists auditory presentation, the response time between the 3 tasks is actually of comparable magnitude.

Regions conjointly activated and conjointly asymmetrical during production, listening and reading of Word-Lists

First observation is that there were 14 hROIs conjointly activated in the left hemisphere and leftward asymmetrical while there were 7 hROIs conjointly activated in the right hemisphere and rightward asymmetrical demonstrating the left hemisphere dominance of brain areas dedicated to Word-List processing (Figure 2, Table 1).

**Table 1.**
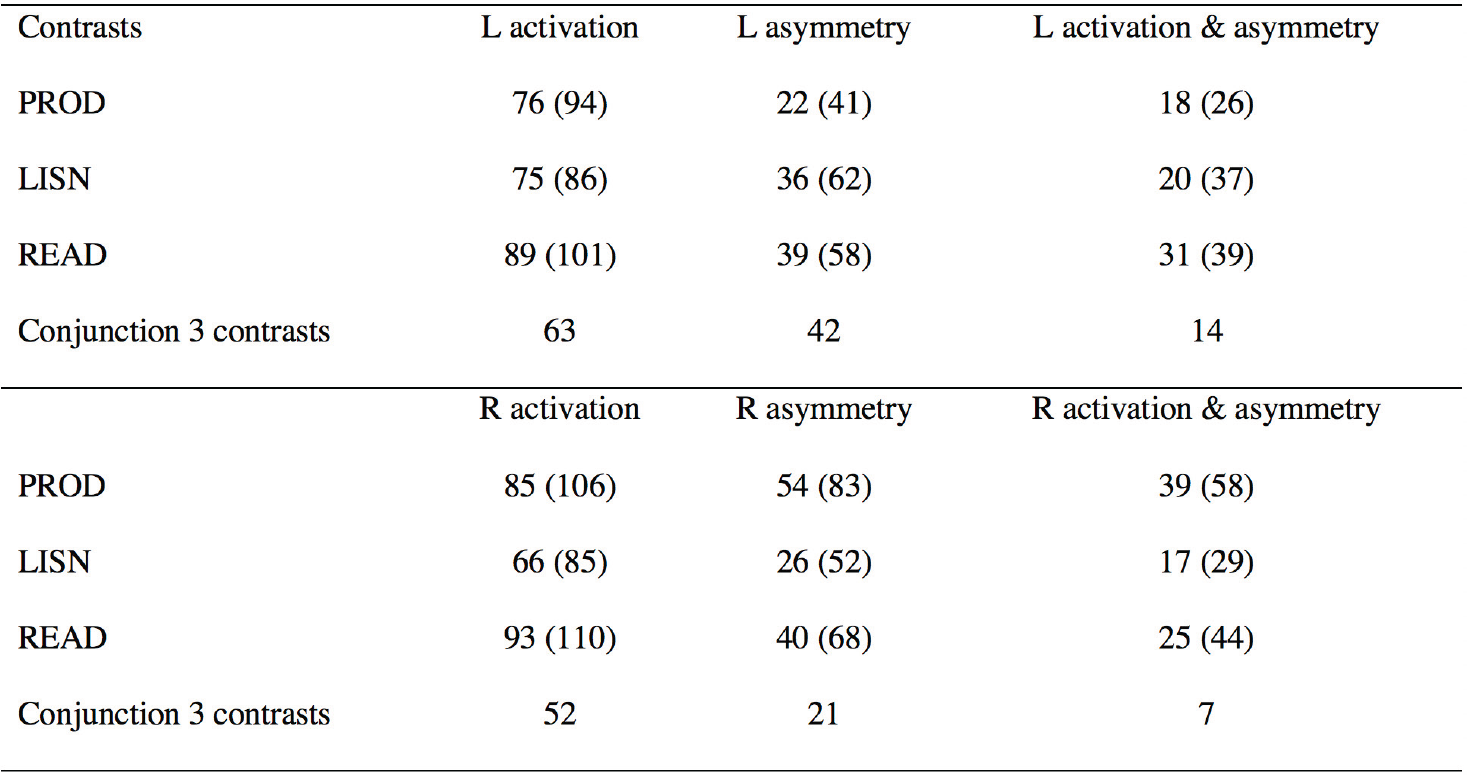
Number of hROIS identified at each selection step. a. Number of hROIs having significant left activation, leftward asymmetry or conjunction of activation and asymmetry for the 3 Word-List tasks. b. Number of hROIs having significant right activation, rightward asymmetry or conjunction of activation and asymmetry for the 3 Word-List tasks. For a given contrast or asymmetry and for a given hemisphere, the statistical threshold was set at 0.00027 (Bonferroni correction for 184 hROIs). For the conjunction between activation and asymmetry the statistical threshold was set at 0.016 (Bonferroni correction for the conjunction of two contrasts) and for the conjunction between 3 tasks, the statistical threshold was set at 0.05 (Bonferroni correction the conjunction of 3 tasks). The number of regions at a non-corrected threshold of 0.05 is given in brackets.

**Fig. 2.**
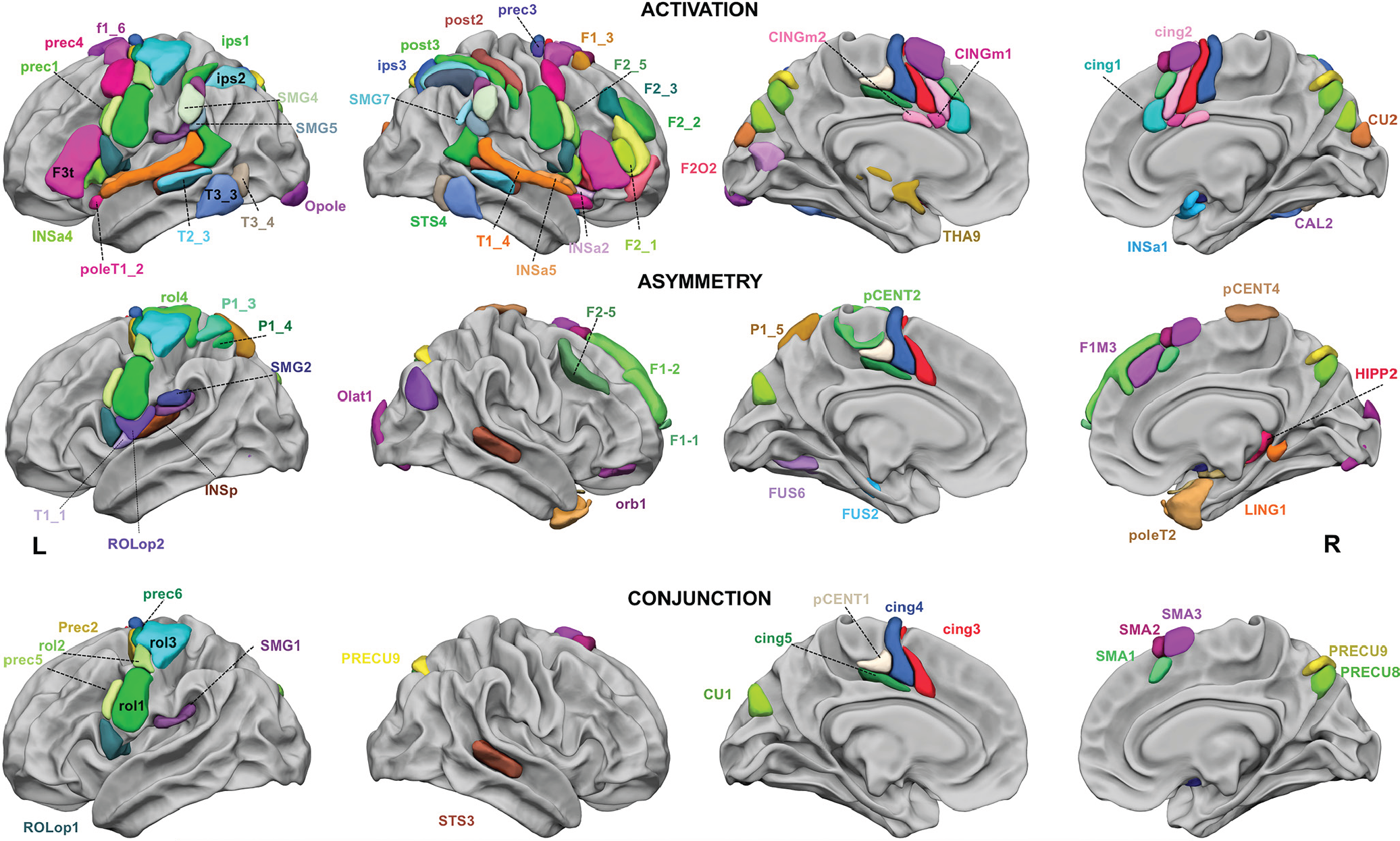
Regions of the AICHA atlas significantly activated in the 3 tasks (top row), significantly asymmetrical in the 3 tasks (middle row), and significantly conjointly activated and conjointly asymmetrical in the 3 tasks (bottom row). The hROIs are projected on the white matter surface of the BIL&GIN template with Surf Ice (https://www.nitrc.org/projects/surfice/) software. Leftward asymmetrical as well as conjointly leftward activated and conjointly asymmetrical hROIS are presented on the left hemisphere and rightward asymmetrical hROIS are presented on the right hemisphere. Selection of hROIs considered as activated or asymmetrical per hemisphere by task was done according to a p-value < 0.05, the statistical threshold applied for conjunction of asymmetry and activation for a given task was p < 0.0025 for each hemisphere and the threshold set for the 3 tasks conjunction was p < 6.25×10^−6^.

#### Left hemisphere

The conjunction of significant leftward activation in the three contrasts (intersection of PROD, LISN and READ) revealed 50 left hROIs (Figure 2, top row). The conjunction of significant leftward asymmetry in the 3 contrasts evidenced 26 hROIs (Figure 2, middle row) and 14 left hROIs that were conjointly conjointly activated and leftward asymmetrical (p < 1.5 × 10^−8^, Figure 2 bottom row, table 1, and see table 2 for abbreviations and coordinates in MNI space). Most of these hROIs were located in the frontal cortex, including 7 hROIs straddling along the Rolandic sulcus (rol1, rol2, rol3, ROLop1) and precentral sulcus (prec2, prec5, prec6) and 4 located on the dorsal part of the internal surface of the frontal lobe (cing3, cing4, cing5, pCENT1). One hROI was located in the parietal lobe, in the supramarginal gyrus (SMG1). Finally, one hROI was located in the occipital lobe, in the cuneus (CU1). Only one subcortical area was selected, the pallidum (PALL).

**Table 2.**
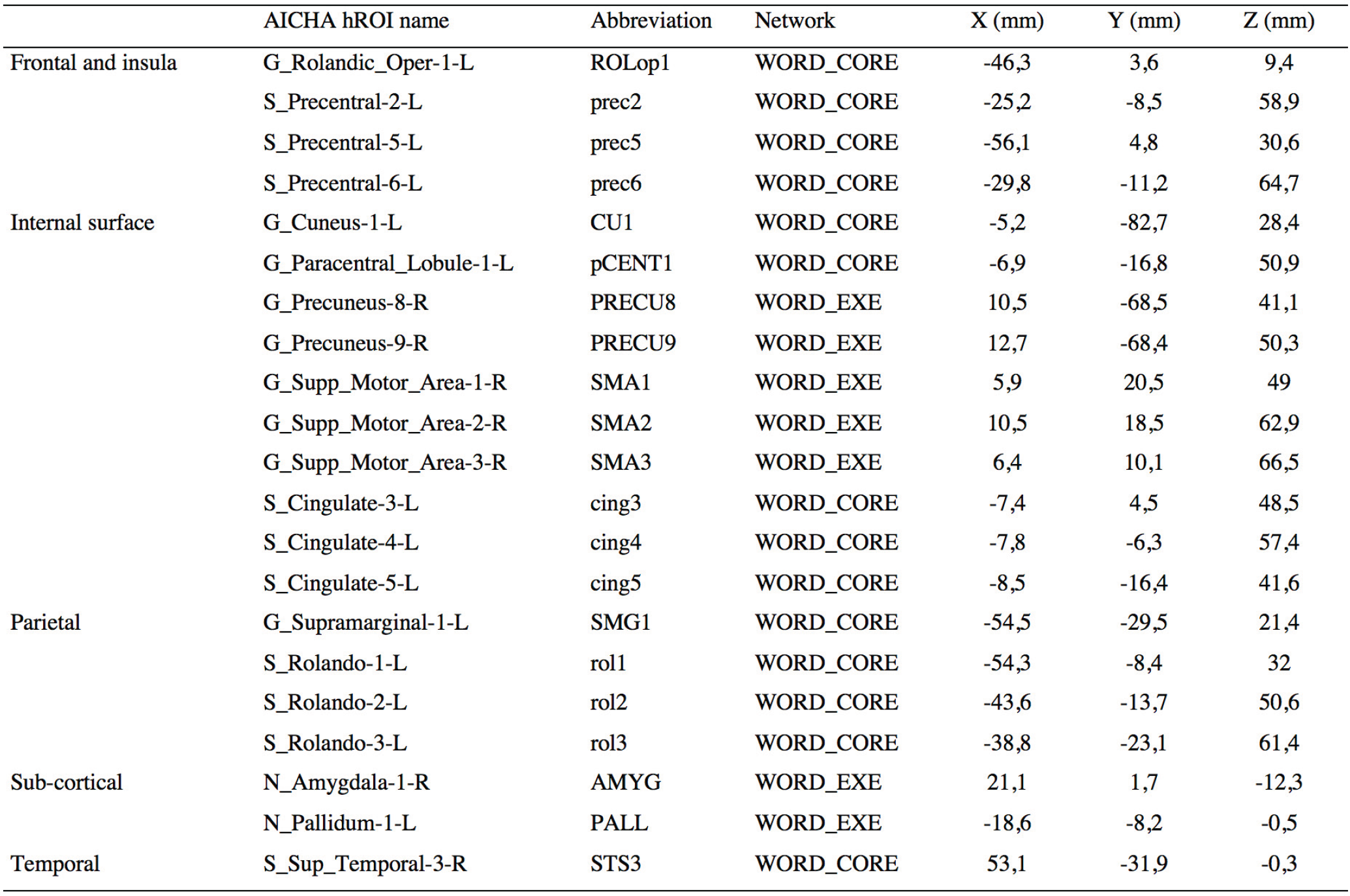
Names and abbreviations of the 21 hROIs selected as either conjointly leftward activated and asymmetrical or conjointly rightward activated and asymmetrical during the 3 Word-List tasks. The network label to which they were clustered and their coordinates in MNI space after SPM12 normalization of the AICHA atlas are also provided.

#### Right hemisphere

The conjunction of significant rightward activation in the three contrasts (intersection of PROD, LISN and READ) revealed 54 hROIs (Figure 2, top row). The conjunction of significant rightward asymmetry evidenced 19 hROIs (Figure 2, middle row) and the conjunction of rightward activation and asymmetry revealed 7 hROIs (Figure 2, bottom row): 3 in the internal surface of the frontal lobe, located anteriorly to those of the left hemisphere (SMA1, SMA2, SMA3), 2 in the precuneus (PRECU8, PRECU9), 1 in the temporal cortex (STS3) straddling the superior temporal sulcus. Only one subcortical area was detected, the amygdala (see table 2 for abbreviations and coordinates in MNI space).

### Activation and asymmetry profiles of the selected hROIs in the 3 tasks

#### Regions of the left hemisphere

There were significant main effects of Task (p<0.0001), ROI (p<0.0001), SIDE (p<0.0001) and all interactions were significant: Task x Side p = 0.002, Task x ROI (p<0.0001), Side x ROI (p<0.0001), Side x task x ROI (p = 0.0002).

Five left regions presented modulation in asymmetry across the different modalities of Word-List processing (Figure 3, table 3). The motor and premotor areas situated in the inferior part of the central sulcus including rol1, and the prec5 (located immediately anterior to rol1) were significantly more asymmetrical and activated during the tasks involving a motor component (PROD and READ) than during LISN. Two adjacent areas located in the mid cingulate cortex (MCC), pCENT1 and cing5, had larger asymmetry during LISN than during READ and PROD, due to a larger right activation during PROD and/or READ than during LISN, while left activations were comparable. Finally, the anterior and dorsal part of the cuneus (CU1), was significantly more asymmetrical and more activated bilaterally during READ than during the 2 other tasks.

**Table 3.** Mean activation in each Word-List contrasts of the 14 hROIs showing joint left activation and left asymmetry during the 3 tasks (abb corresponds to the abbreviation of the AICHA hROI name provided in table 2).

| hROIs abb | PROD |  |  | LISN |  |  | READ |  |  |
| --- | --- | --- | --- | --- | --- | --- | --- | --- | --- |
|  | Mean (SD) | t | p | Mean (SD) | t | p | Mean (SD) | t | p |
| **Left activation** |  |  |  |  |  |  |  |  |  |
| CU1 | 0.24 (0.38) | 7.45 | <.0001 | 0.14 (0.35) | 4.87 | <.0001 | 0.55 (0.71) | 9.32 | <.0001 |
| pCENT1 | 0.14 (0.27) | 6.36 | <.0001 | 0.18 (0.24) | 9.30 | <.0001 | 0.13 (0.34) | 4.65 | <.0001 |
| ROLOp1 | 0.51 (0.27) | 22.93 | <.0001 | 0.24 (0.26) | 10.98 | <.0001 | 0.28 (0.33) | 10.28 | <.0001 |
| SMG1 | 0.41 (0.39) | 12.60 | <.0001 | 0.94 (0.63) | 17.95 | <.0001 | 0.12 (0.49) | 2.98 | 0.0034 |
| PALL | 0.25 (0.19) | 16.09 | <.0001 | 0.11 (0.16) | 8.23 | <.0001 | 0.42 (0.27) | 19.00 | <.0001 |
| cing3 | 0.83 (0.39) | 25.61 | <.0001 | 0.47 (0.32) | 17.81 | <.0001 | 0.69 (0.49) | 16.74 | <.0001 |
| cing4 | 0.52 (0.34) | 18.71 | <.0001 | 0.27 (0.26) | 12.40 | <.0001 | 0.38 (0.43) | 10.46 | <.0001 |
| cing5 | 0.15 (0.26) | 6.88 | <.0001 | 0.15 (0.22) | 8.32 | <.0001 | 0.16 (0.31) | 6.00 | <.0001 |
| prec2 | 0.12 (0.23) | 6.37 | <.0001 | 0.08 (0.21) | 4.48 | <.0001 | 0.24 (0.39) | 7.30 | <.0001 |
| prec5 | 0.76 (0.49) | 18.67 | <.0001 | 0.40 (0.36) | 13.46 | <.0001 | 0.91 (0.54) | 20.09 | <.0001 |
| prec6 | 0.20 (0.36) | 6.54 | <.0001 | 0.19 (0.32) | 7.26 | <.0001 | 0.24 (0.49) | 5.80 | <.0001 |
| rol1 | 0.66 (0.51) | 15.45 | <.0001 | 0.26 (0.29) | 11.01 | <.0001 | 0.54 (0.50) | 13.03 | <.0001 |
| rol2 | 0.61 (0.43) | 16.85 | <.0001 | 0.40 (0.33) | 14.78 | <.0001 | 0.72 (0.52) | 16.55 | <.0001 |
| rol3 | 0.21 (0.31) | 7.91 | <.0001 | 0.17 (0.32) | 6.53 | <.0001 | 0.28 (0.43) | 7.70 | <.0001 |
| **Asymmetry (Left – Right)** |  |  |  |  |  |  |  |  |  |
| CU1 | 0.04 (0.19) | 2.75 | 0.0068 | 0.05 (0.17) | 3.70 | 0.0003 | 0.15 (0.34) | 5.40 | <.0001 |
| pCENT1 | 0.13 (0.12) | 12.8 | <.0001 | 0.20 (0.18) | 13.88 | <.0001 | 0.15 (0.18) | 9.89 | <.0001 |
| ROLOp1 | 0.07 (0.23) | 3.62 | 0.0004 | 0.10 (0.21) | 5.57 | <.0001 | 0.11 (0.27) | 4.86 | <.0001 |
| SMG1 | 0.26 (0.34) | 9.23 | <.0001 | 0.25 (0.69) | 4.37 | <.0001 | 0.35 (0.43) | 9.69 | <.0001 |
| PALL | 0.03 (0.13) | 2.98 | 0.0034 | 0.04 (0.12) | 3.36 | 0.0010 | 0.03 (0.17) | 2.02 | 0.0451 |
| cing3 | 0.07 (0.26) | 3.36 | 0.0010 | 0.10 (0.20) | 5.72 | <.0001 | 0.11 (0.30) | 4.32 | <.0001 |
| cing4 | 0.09 (0.21) | 5.28 | <.0001 | 0.11 (0.16) | 7.77 | <.0001 | 0.13 (0.22) | 6.97 | <.0001 |
| cing5 | 0.05 (0.14) | 4.34 | <.0001 | 0.15 (0.16) | 11.34 | <.0001 | 0.10 (0.18) | 6.70 | <.0001 |
| prec2 | 0.08 (0.20) | 5.14 | <.0001 | 0.12 (0.20) | 6.92 | <.0001 | 0.12 (0.30) | 4.54 | <.0001 |
| prec5 | 0.37 (0.53) | 8.26 | <.0001 | 0.14 (0.42) | 3.93 | 0.0001 | 0.48 (0.51) | 11.42 | <.0001 |
| prec6 | 0.15 (0.33) | 5.45 | <.0001 | 0.18 (0.35) | 6.12 | <.0001 | 0.16 (0.43) | 4.37 | <.0001 |
| rol1 | 0.27 (0.29) | 11.29 | <.0001 | 0.08 (0.24) | 4.20 | <.0001 | 0.25 (0.29) | 10.49 | <.0001 |
| rol2 | 0.43 (0.35) | 14.69 | <.0001 | 0.37 (0.36) | 12.32 | <.0001 | 0.44 (0.43) | 12.25 | <.0001 |
| rol3 | 0.21 (0.29) | 8.42 | <.0001 | 0.18 (5.88) | 0.37 | <.0001 | 0.20 (0.40) | 5.88 | <.0001 |

**Fig. 3.**
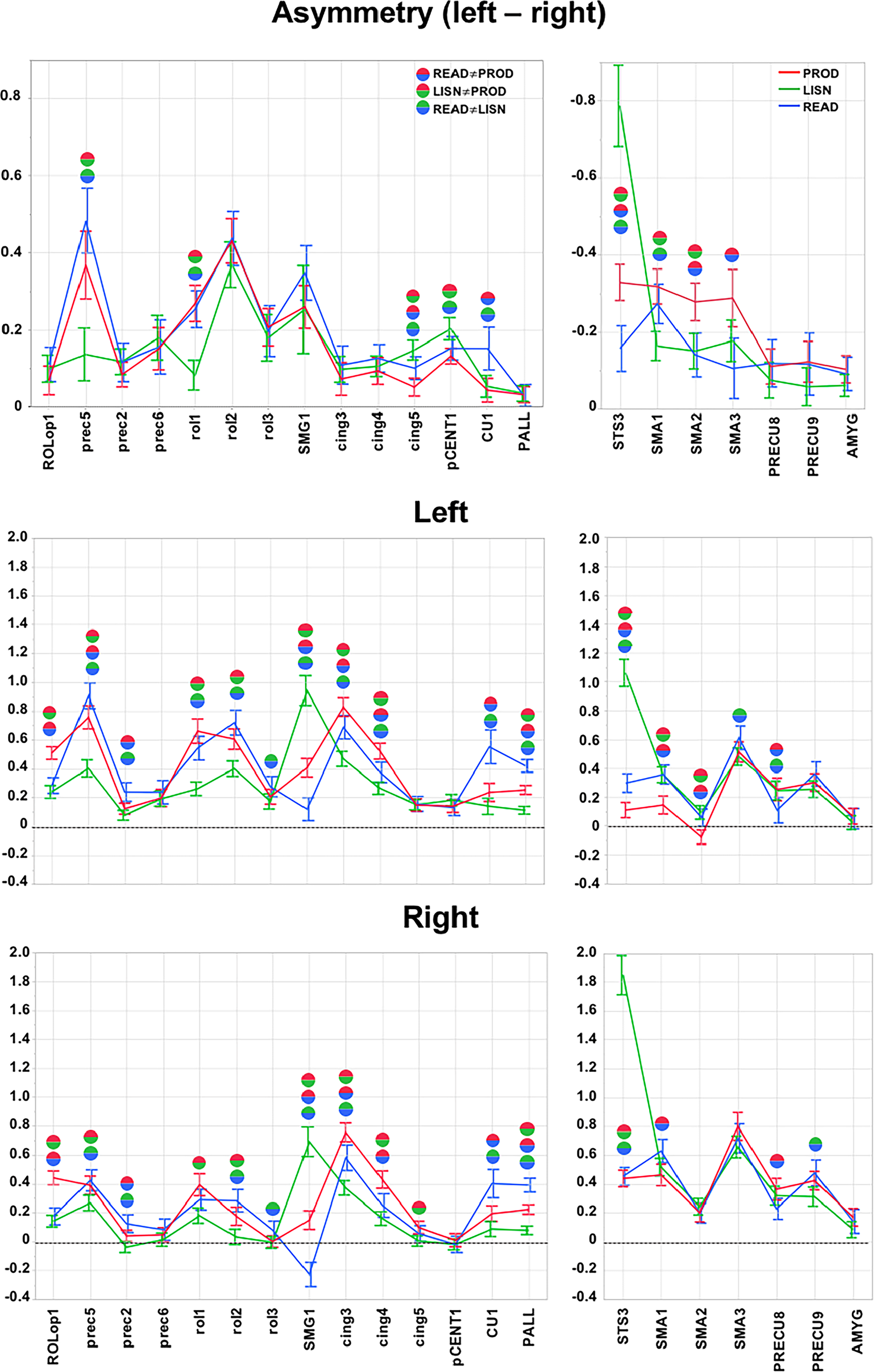
Activity and asymmetry measured in the 21 hROIs selected at step 1. The 14hROIs selected as leftward asymmetrical and activated are displayed on the left column and the 7 hROIs selected as rightward and asymmetrical are displayed on the right column. Top row are the asymmetry values in each task (red corresponds to PROD, green to LISN, and blue to READ, note that the scale of the right hROIs has been inverted to facilitate the comparison with left regions) and significant differences (Bonferroni corrected for the number of hROIs) across tasks are indicated by spheres (blue-red sphere corresponds to a significant difference between PROD and READ, red-green sphere corresponds to a significant difference between PROD and LISN, and blue-green sphere corresponds to a significant difference between LISN and READ. Middle row are the left values of activity in the left hemisphere of the same set of regions, and bottom row the activity in the right hemisphere. Abbreviations correspond to those provided in table 2.

A second set of motor regions showed little modulation of their asymmetry by the task modality. A first set corresponding to cing3 and cing4 showed no difference in asymmetry across tasks but larger activity during PROD and READ, which present a stronger motor component than during LIST. A second region, ROLop1, situated in the Rolandic operculum area, presented no difference in asymmetry across task but revealed larger activity in PROD as compared to the two other tasks. A third set, including prec2 and prec6 located immediately anteriorly to rol3, was also characterized by little inter-tasks differences in activation.

Finally, most of the regions selected in the left hemisphere exhibited bilateral activations larger during the tasks having a stronger motor component (PROD and READ) than during LISN (Figure 3). Note that SMG1 bilateral activity was strongly increased by the auditory modality, with stronger activation during LISN as compared to the 2 other tasks.

#### Regions of the right hemisphere

There were significant main effect of Task (p<0.0001), ROI (p<0.0001), SIDE (p<0.0001) and all interactions were significant: Task x Side p = 0.0031, Task x ROI (p<0.0001), Side x ROI (p<0.0001), Side x task x ROI (p <0.0001).

As for the left regions, all the right hROIs showed bilateral activation except SMA2 which was deactivated on the left during PROD (Figure 3, table 4).

**Table 4.** Mean activation and asymmetry in each Word-List contrasts of the 7 hROIs showing right activation and right asymmetry during the 3 tasks (abb corresponds to the abbreviation of the AICHA hROI name provided in table 2).

| hROIs abb | PROD |  |  | LISN |  |  | READ |  |  |
| --- | --- | --- | --- | --- | --- | --- | --- | --- | --- |
|  | Mean (SD) | t | p | Mean (SD) | t | p | Mean (SD) | t | p |
| **Left activation** |  |  |  |  |  |  |  |  |  |
| PRECU8 | 0.36 (0.46) | 9.48 | <.0001 | 0.32 (0.39) | 9.85 | <.0001 | 0.23 (0.44) | 6.18 | <.0001 |
| PRECU9 | 0.42 (0.39) | 13.04 | <.0001 | 0.31 (0.4) | 9.34 | <.0001 | 0.48 (0.55) | 10.42 | <.0001 |
| SMA1 | 0.46 (0.45) | 12.24 | <.0001 | 0.52 (0.37) | 16.83 | <.0001 | 0.63 (0.49) | 15.36 | <.0001 |
| SMA2 | 0.20 (0.38) | 6.34 | <.0001 | 0.24 (0.35) | 8.35 | <.0001 | 0.20 (0.42) | 5.70 | <.0001 |
| SMA3 | 0.80 (0.58) | 16.43 | <.0001 | 0.66 (0.46) | 17.26 | <.0001 | 0.72 (0.61) | 14.04 | <.0001 |
| AMYG | 0.17 (0.37) | 5.48 | <.0001 | 0.08 (0.34) | 2.97 | 0.0035 | 0.14 (0.51) | 3.34 | 0.0011 |
| STS3 | 0.49 (0.35) | 15.05 | <.0001 | 1.85 (0.82) | 27.06 | <.0001 | 0.45 (0.37) | 14.70 | <.0001 |
| **Asymmetry (Left – Right)** |  |  |  |  |  |  |  |  |  |
| PRECU8 | -0.11 (0.28) | -4.77 | <.0001 | -0.07 (0.28) | -3.18 | 0.0018 | -0.12 (0.37) | -3.82 | 0.0002 |
| PRECU9 | -0.12 (0.32) | -4.55 | <.0001 | -0.06 (0.30) | -2.30 | 0.0226 | -0.12 (0.49) | -2.85 | 0.0051 |
| SMA1 | -0.32 (0.28) | -13.85 | <.0001 | -0.16 (0.23) | -8.56 | <.0001 | -0.27 (0.31) | -10.51 | <.0001 |
| SMA2 | -0.28 (0.30) | -11.28 | <.0001 | -0.15 (0.28) | -6.43 | <.0001 | -0.14 (0.35) | -4.86 | <.0001 |
| SMA3 | -0.29 (0.45) | -7.64 | <.0001 | -0.18 (0.33) | -6.48 | <.0001 | -0.11 (0.48) | -2.65 | 0.0089 |
| AMYG | -0.10 (0.22) | -5.74 | <.0001 | -0.06 (0.17) | -4.22 | <.0001 | -0.09 (0.26) | -4.13 | <.0001 |
| STS3 | -0.33 (0.28) | -13.85 | <.0001 | -0.79 (0.64) | -14.73 | <.0001 | -0.16 (0.36) | -5.22 | <.0001 |

One region had a very different profile than all the others, STS3, located at the mid-third of the STS. STS3 presented significantly larger rightward asymmetry during LISN, then PROD and READ. STS3 was also significantly more activated bilaterally during LISN than during the two other tasks (Figure 3).

Three hROIS of the internal surface of the frontal lobe, labelled SMA, located at the dorsal face and anterior part of the medial frontal gyrus, also showed modulation of their asymmetry by the modality of the Word-List tasks. SMA1 was significantly more rightward asymmetrical during PROD and READ than during LISN whereas SMA2 and SMA3 were significantly more asymmetrical during PROD than during the 2 other tasks.

On the opposite, the PRECU8, PRECU9 and AMYG did not show any variation in asymmetry with the task modality.

Apart from STS3, profiles of activations were comparable across tasks. However, there was smaller left activation in SMA1, SMA2 and SMA3 during PROD (with a deactivation for SMA3), and smaller left activation during READi in PRECU8. On the right, activation was smaller during READi in PRECU8 and PRECU9 as compared to PROD and LISN respectively, and in SMA1 during PROD as compared to READi.

### Part 2. Intra- and Inter-hemispheric connectivity at rest

## Method

### Participants

A subset of 138 participants (mean age 27.3 years (SD = 6.4 years, 68 women) also completed a resting-state fMRI (rs-fMRI) acquisition lasting 8 minutes. Note that this resting-state acquisition was performed on average 9 months (SD = 9.6 months) before the language task acquisition in all but 5 cases. In these 5 cases, the resting-state acquisition occurred approximately one year after the language session (range [11.2 - 13.8] months).

### Resting-stateimageacquisition(rs-fMRI) and processing

Spontaneous brain activity was monitored for 8 min (240 volumes) using the same imaging sequence (T2*-weighted echo-planar images) as that used for the language tasks. Immediately prior to rs-fMRI scanning, the participants were instructed to “keep their eyes closed, to relax, to refrain from moving, to stay awake and to let their thoughts come and go” (Mazoyer et al. 2016). After identical pre-processing than that used for task-induced fMRI acquisitions, time series white matter and cerebrospinal fluid (individual average time series of voxels that belonged to each tissue class) and temporal linear trends were removed from the rs-fMRI data series using regression analysis. Additionally, rs-fMRI data were temporally filtered using a least squares linear-phase finite impulse response filter design bandpass (0.01– 0.1 Hz).

For each of the 138 participants that completed resting-state acquisition and for each of the same 185 homotopic ROIs, an individual BOLD rs-fMRI time series was computed by averaging the BOLD fMRI time series of all voxels located within the hROI volume.

From the BOLD fMRI time series of hROIs, we computed the correlations of each hROI pair of the 21 selected hROIs in each participant, we then averaged the correlations among pairs of hROIs across the 138 individuals resulting in a matrix of intrinsic connectivity for the whole population.

### Resting-state image analysis: characterizing networks within the 21 selected hROIs

Using the same methodology as in Labache et al (2019), we applied an agglomerative hierarchical cluster analysis method on the intrinsic connectivity matrix to identify the different networks supporting the organization across the 21 hROIs. We tested the reliability of the identified networks using a multiscale bootstrap resampling method that provides us with an AU p value that represents the stability of networks based on the average matrix of intrinsic connectivity.

Finally, we calculated the average functional intrinsic correlations between the identified networks. The significance of these correlations compared to 0 was assessed using a non-parametric sign test at a significance level of 0.05 (Bonferroni correction for the number of network pairs).

## Results

### Identification of networks based on the resting-state connectivity of the 21 hROIs conjointly leftward activated and asymmetrical or rightward activated and asymmetrical during the 3 Word-List tasks

Hierarchical clustering analysis revealed 2 networks from the selected set of 21 hROIs (Figure 4), one we la-belled WORD_CORE composed of 13 left hROIs and one right hROI and the other one we labelled WORD_ EXE composed of 1 left area and of 6 right hROIs.

**Fig. 4.**
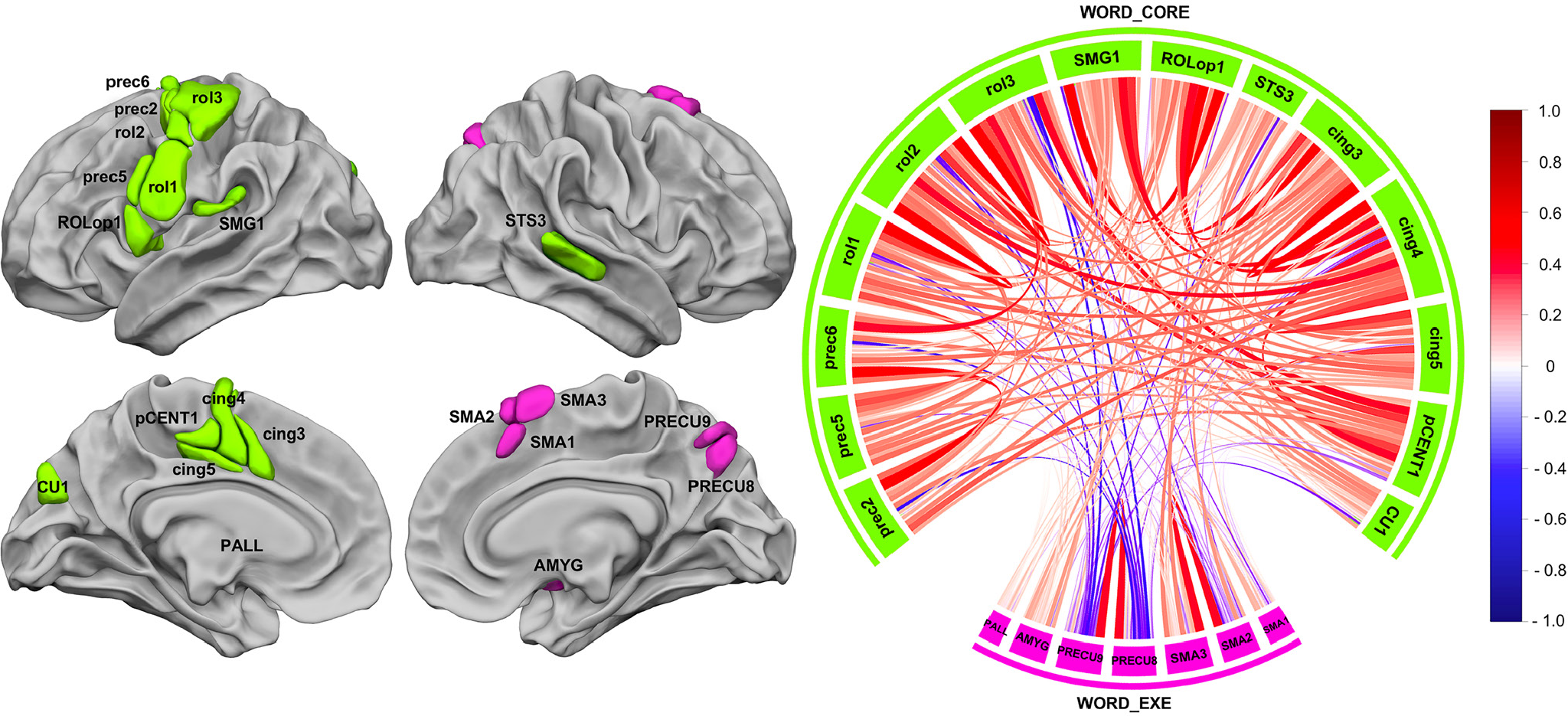
Intra- and inter-hemispheric correlation at rest across the 21 hROIs selected as either conjointly leftward activated and asymmetrical or conjointly rightward activated and asymmetrical during the 3 language tasks. The left motor areas and the right STS are strongly and positively correlated and they constitute the WORD-CORE network (green) that is not significantly anticorrelated with the WORD_EXE network, which is made of the right precuneus and SMA regions and the left pallidum (pink). The chord diagram was produced by R with the package “circlize” (Gu Z. et al, 2014).

#### WORD_CORE network

WORD_CORE (Figure 4, green) was composed of 13 hROIs hosting motor and premotor areas of the left hemisphere gathered along the Rolandic sulcus (rol1, rol2, rol3), precentral sulcus (prec2, prec5), Rolandic operculum (ROLop1), motor and premotor areas of the medial surface (pCENT1, cing3, cing4, cing5) and SMG1 of the parietal lobe. Importantly WORD_CORE network is also an inter-hemispheric network, since it comprised the right STS3 in addition to the 13 left hROIs. We named this network WORD_CORE because it included essential phonological processing regions, as further described below. WORD_CORE was the largest network in terms of volume (53136 mm^3^), as it was 4.05 times larger than WORD_EXE (13104 mm^3^).

ROLop1 appeared to be a very important hROI for communication within this network since it was among the top 5% of the strongest correlations among non-contiguous hROIs (ROLop1-SMG1: r=0.62, ROLop1-cing3: r=0.59, ROLop1-cing4: r=0.52). It is interesting to note that the SMG1-cing4 correlation (r=0.53) was also part of the 5% highest correlation underlying a very strong antero-posterior tripartite connection between ROLop1, SMG1 and cing4.

#### WORD_EXE network

The second network was constituted of 7 hROIs : 6 right hROIs including the 3 superior motor areas of the internal frontal lobe surface (SMA1, SMA2 and SMA3, which is SMA proper), the posterior part of the precuneus (PRECU8 and PRECU9) and AMYG. In addition to these right areas, the left PALL was also part of the network. We labelled it WORD_EXE because it included regions involved in the achievement of the low-level reference task (button click).

Note that the AU p values provided by the multiscale bootstrap resampling method showed that both networks were reliable at 81%.

#### Temporal correlation across networks and significance

The chord diagram shown in Figure 4 describes the average correlations between each pair of hROIs in the two networks. A non-significant negative mean intrinsic correlation was found between WORD_CORE and WORD_ EXE (R = −0.04; 55.07% of the participants showed a negative correlation, p = 0.13).

### Summary of the results of the whole study

The analysis of the connectivity at rest across the 21 hROIs common to the production, listening and reading of over learnt lists of words, made it possible to identify a set of two networks. A large WORD_CORE network made of plurimodal areas was revealed, including on the left hemisphere, premotor and motor regions, which were more activated during PROD and READ than during LISN, an auditory area situated in the anterior part of the left SMG, which was more activated during LISN than during PROD and READ, and a visual region, CU1, which was more activated and more asymmetrical during READ as compared to the 2 other tasks. Importantly the strongest correlations at rest between these 21 hROIS were observed across anterior and posterior areas (action-perception), namely Rolop1, and cing4 with SMG1. On the right hemisphere, the WORD_CORE network included the STS3, located on the mid-third of the sulcus, which was significantly more activated and asymmetrical during LISN than during PROD and READ. Note that a second network (WORD_ EXE), made of right SMA1 and SMA2 (located at the pre-SMA), SMA3 (SMA proper), the precuneus, as well as the left pallidum (PALL), was identified and was not correlated with the WORD_CORE network. Areas composing this WORD_EXE are mainly involved in extra linguistic, executive processes and attentional systems recruited to manage task completion.

## Discussion

Here we evidence a large scale network of areas commonly shared by the production, listening and reading of lists of words, which not only includes articulatory and auditory areas but also a region of the visual cortex, the cuneus.

Though more recruited during the reading task, this region, considered to be a component of accurate phonological awareness (Bolger et al. 2008), is also at play during word-list articulation and listening. During development, speech perception and production, engaging auditory and motor (articulation) modalities, are first linked together. The later development of reading skills, engaging the visual modality, is elaborated upon these 2 components. The present results evidence a brain organization in adults that is a reflection of the whole developmental and learning processes of language, where action and perception circuits are interdependent and organised in networks, among which a trace of the learning modality is still present in the brain.

We will first discuss the WORD_CORE network from the left motor and premotor areas action side, up to the involvement of the audio-motor loop extending to the phonological loop and the right STS.

### WORD_CORE network underpinning supramodal word-list processing

#### Left motor and premotor areas: from the speech effectors areas to the hand area

On the action side, results have revealed 7 areas straddling along the Rolandic sulcus and precentral gyrus and 4 located the internal surface of the frontal lobe which were modulated by the mere nature of the task depending on the modality.

Regions having the strongest motor involvement were located along the Rolandic sulcus and included primary motor areas that showed huge activations in PROD and READ. This is in accordance with Penfield’s cortical stimulation studies, which have provided the first functional support for the existence of somatotopy within the orofacial region (Penfield and Roberts, 1959). In fact, rol1 and adjacent rol2 not only matches the area involved in speech production as described by Wilson (Wilson et al., 2004) but also areas involved in mouth, larynx, tongue, jaw and lips movements, reported by several studies (Brown et al., 2009, 2008; Fox et al., 2001; Grabski et al., 2012; Wilson et al., 2004) (Figure 5). More precisely, along the dorsal-to-ventral orientation of rol1, the representation for speech listening, mouth, larynx, lips, jaw, tongue, larynx clearly resembles the somatotopic organization of speech effectors proposed by Conant and collaborators (Conant et al., 2014). Moreover, the stronger asymmetry in this area during PROD and READ is consistent with the fact that these two tasks involve covert articulation or subvocalization. Prec5, tightly joining rol1 along with cing3, was characterized by very strong BOLD increase during READ as compared to both PROD and LISN, suggesting a motor preparation activity (Dietz et al., 2005). On the internal surface, cing3 and cing4, partly overlapping the SMA tongue area, and the anterior part of the mid cingulate cortex situated at the tongue cingulate motor area according to Amiez et al (2014), also presented a strong motor component as revealed by their increased activation in PROD and READ as compared to LISN. Note that the left PALL was more activated during READ as compared to the 2 other tasks, which is in accordance with results from a meta-analysis revealing that adult readers recruited a large network including the left PALL as compared to children (Paulesu et al, 2014). Importantly, the same region has been shown to be at play in the audio-motor adjustments during auditory feedback control of speech, confirming its involvement in plurimodal modulation (Tourville et al. 2008).

**Fig. 5.**
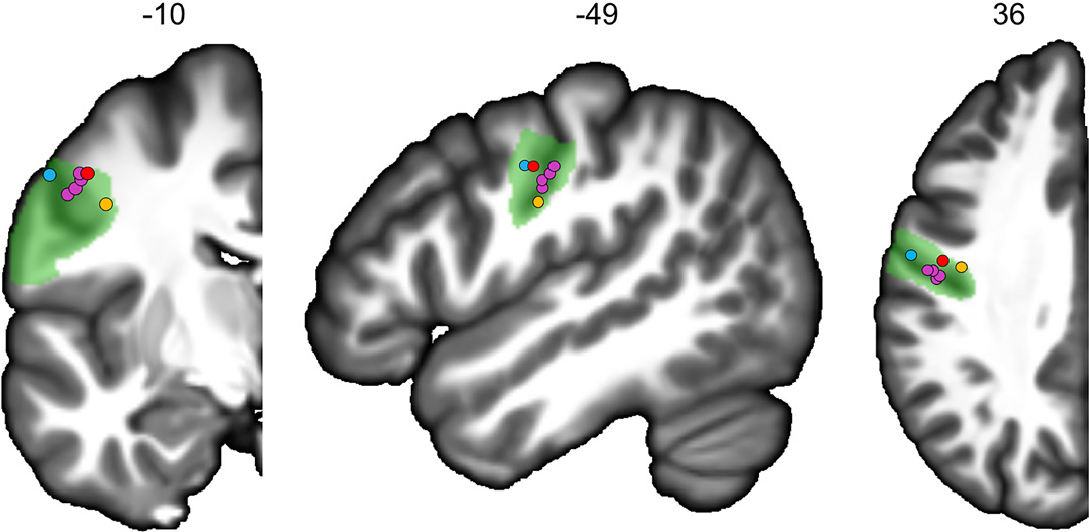
Locations of the activation picks of 4 studies on the left hemisphere coronal, sagittal and axial slices from the BIL&GIN display template; the hROI numbers correspond to the x, y and z-axis in the MNI space. In green, representation of rol1. Larynx areas are in blue (Brown et al., 2009) and orange (Brown et al., 2008). In red, mouth area (Fox et al., 2001). In purple, lips, tongue, jaw and vowels areas (Grabski et al., 2012).

On the most ventral part of the Rolandic sulcus, ROLop1 presented a greater involvement in PROD. This region has been reported to be activated by overt and covert articulation (Heim et al., 2002; Herbster et al., 1997; Price et al., 1996), to be involved in phonological rehearsal (Veroude et al., 2010), in silent recitation of the months of the year (Wildgruber et al., 1996), and its lesion has been associated with articulatory disorders (Tonkonogy and Goodglass, 1981), which is in accordance with the motor component revealed in the present study. Its lower activation as compared to rol1 and prec5 is due to the fact that this region is a secondary motor area. It could also be put forward that ROLop1 activation could implement a simulation of phoneme production according to the model by Wilson & Iacoboni (Wilson and Iacoboni, 2006), which stipulates that the prediction of the acoustic consequences of phonemes production would be compared in the superior temporal cortex with the acoustic analysis of the heard speech sounds. In the present study, we cannot disentangle between an actual motor component related to subvocalization from a simulation, both being potentially at play depending on the task, i.e., simulation during the listening task and vocalization during the production task as well as a very likely vocalization aspect during the reading task. Moreover, among the WORD_CORE network, intrinsic connectivity at rest has revealed that ROLop1 is an essential hROI for communication within this network, being particularly correlated with not only mid cingulate cortex tongue areas, cing3 and cing4, but also with SMG1.

On the dorsal part, a last set of areas, prec2 and prec3 premotor areas situated along the precentral sulcus and tightly joining rol3, which is at the location of the hand-motor area (Mellet 2016), were conjointly activated during the 3 tasks, and did not show strong differences among tasks. On the internal surface, pCENT1 and cing5, located in the mid cingulate cortex have also been revealed to be involved in the hand representation (Amiez et al, 2014). Given their functional location, one explanation could be put forward concerning the language ontogenesis observations stipulating that the motor control of the vocal tract (speech articulators) and the motor control of the hand develop in cooperation, arguing for a hand-mouth gestural basis for language (Iversen, 2010). An alternative explanation would be in relation with the motor response provided by the subjects at the end of each Word-List task, however, as the central cross change detection likely cancels out most of these non-specific hand-motor activations as well as the lack of somatosensory activation does not seem to support the latter hypothesis.

#### The audio-motor loop is involved in READ and extends to the phonological loop

Results from the intrinsic connectivity evidenced a strong correlation between SMG1 and cing4, underlying a very strong antero-posterior tripartite connection between RO-Lop1, SMG1 and cing4. So, there does exist a group of hROIs common to the 3 Word-List tasks, with a leftward involvement of brain areas specifically involved in word processing which were found to be strongly and positively synchronised at rest, constituting a network that included the more anterior left SMG hROI, namely SMG1 as well as frontal and cingulate motor and premotor areas. These frontal and temporal regions, connected through the arcuate fasciculus tract (Catani et al., 2005), appear to be composed of hROIs involved in plurimodal representation of speech sound processing, the so-called audio-motor loop (Vigneau et al., 2006). The left areas of the audio motor network correspond at large to the neural bases of a perception-action cycle for speech sounds conjointly recruited during the three Word-List tasks. Such a model, based on the link between speech perception and production, posits that articulatory gestures are the central motor units on which both speech perception and production develop and act according to the motor theory of speech (Liberman and Whalen, 2000). More recently, neurobiological theories of speech perception have proposed a more dynamic and integrative model in which language processing would rely on action-perception circuits distributed across auditory and motor systems (Pulvermüller and Fadiga, 2016, 2010). The fact that, within this network, conjointly activated and asymmetrical areas, which hosted mostly motor and premotor areas, were also activated during LISN, though at a lower intensity and with a lower asymmetry, together with the recruitment of the SMG1 in the production task, favours the theory supporting an involvement of action-perception circuits whatever the Word-List task (Pulvermüller and Fadiga, 2016, 2010). Actually, it is interesting to observe that READ also appears to recruit this action-perception circuit, revealing a significant activation in rol1 as compared to LISN. Interestingly, this circuit, posited to subserve the mapping between sensory and motor phonological codes, is also engaged in reading (Danelli et al, 2015; Malins et al, 2016). The action-perception circuit recruited in the present study for READ also reflects the fact that the participants were instructed to engage their attention in the reading of each word and covertly articulate the lists of words in a slow speech rate. The fact that ROLop1 activation in the present study is similar for READ and LISN favours the recruitment of the action-perception circuit reflecting that while reading these over-learnt words, subjects tended to subvocalize them, which is supported by the gradient of significant activation in rol1 (production > reading > listening).

Within the large perception-action model of Fuster (2001, 2014), the literature has identified a set of areas that are considered as the neural support of the phonological working memory loop postulated by Baddeley (Baddeley et al., 1998). Actually, dealing with Word-List automatically engage working memory processes and that component is common to the 3 word tasks. On the perception side, one cluster (SMG1) was more activated and asymmetrical in LISN as compared to READ and PROD. Interestingly, the SMG1 situated on the posterior part of the planum temporale, corresponding in the literature to the Sylvian-parietal-temporal area or Spt (Buchsbaum et al., 2011; Yue et al., 2018), has been described as a sensory-motor integration area for the vocal tract. Area Spt would be part of the phonological loop described by Baddeley (Baddeley et al., 1998), in which the content of the phonological store can be kept active via articulatory rehearsal (Buchsbaum et al., 2011). More precisely, Spt area has been assumed to have a storage function (Martin, 2005; Smith and Jonides, 1998) and to play an important role in the short-term storage of phonological representations by serving as a phonological buffer (Yue et al., 2018). The activity of area Spt would be correlated with frontal speech production areas even if their precise locations differ according to the studies: the pars opercularis according to Buchsbaum et al, (2001) and Poldrack et al, (Poldrack et al., 1999) and the dorsal part of the pars triangularis of the inferior frontal gyrus according to the meta-analysis by Vigneau et al (2006). It is noteworthy that F3t was bilaterally activated during the 3 tasks in the present study (Figure 2, top row), this bilateral involvement impeding a conjoint activation and asymmetry. Together with the F3t activation, on the action side, prec5 was found to be conjointly activated and left asymmetrical by the 3 Word-List tasks, with a gradient of activation ranging from more activation for READ and PROD and less activation for LISN. This premotor area, prec5, has been proposed to make up a subvocal rehearsal system (Chein and Fiez, 2001) and/or to support executive processes allowing for maintaining the content of verbal working memory (Smith and Jonides, 1998), which is in accordance with the highest activation during READ and PROD. Furthermore, a single rTMS intervention targeting either the left SMG or the left posterior part of the inferior frontal gyrus, which are considered as phonological nodes, was already sufficient to disrupt phonological decisions, providing further support that both regions contribute to efficient phonological decisions, and more particularly, subvocal articulation (Hartwigsen et al., 2016).

#### The right STS3 involved in prosodic integration is also activated during PROD and READ and strongly connected with motor areas within WORD_CORE network

The rightward conjoint activation of the STS3 as well as its rightward asymmetry, though more activated in LISN and less activated in READ, could be accounted for by a rightward preference for non-linguistic information such as tonal prosody (Belin et al., 2002). More particularly, the right STS3, which is located on the mid-part of the STS, closely matches the activation peak showing rightward asymmetry described by Beaucousin and others (2007), overlapping the posterior spot of the Human Voice Area (pHVA, Pernet el al. 2015) and also corresponds to the posterior voice area (pHVA) described by Bodin et al. (2018). pHVA is located on a sulcal pit corresponding to the place of the larger sulcal depth. In addition, the rightward asymmetry of this sulcal depth is specific to Humans and exists whatever the age (infants, children or adults), (Leroy et al, 2015). The present study brings elements on the precise role of this region, which is still not fully understood. We propose that the metrics of lists of the words, resembling a reciting, are processed in this area, which is supported by the greater activation during LISN. This is in accordance with the auditory material proposed to the participants, the list being spoken along with the specific prosody of a list. Moreover, this region has been revealed to be more activated during a prosodic task as compared to a phonetic task (Sammler, 2015). During speech, words present non-verbal prosodic information, which is intertwined with the verbal information (Kotz and Paulmann, 2007; Pell and Kotz, 2011). Furthermore, prosodic and verbal cues in speech differ in their spectro-temporal properties: prosodic cues consisting of slow spectral changes (over more than 200ms) and phonemic cues consisting of fast spectral changes (less than 50ms). The right hemisphere has been suggested to be more sensitive to fine spectral details over relatively long time scales and the left hemisphere more sensitive to brief spectral changes (Boemio et al., 2005; Poeppel et al., 2008; Zatorre and Belin, 2001).

The link between the STS3 (corresponding to pHVA), which is a prosodic integrative area, with the left SMG and rol1 instantiated in the resting-state connectivityapproach reflects the intertwining between prosodic and phonemic information. The present results are supported by a recent fMRI/ERP study which has revealed that activity in the left SMG together with the central sulcus area occurs earlier than in the left superior temporal cortex during the phonological processing of ambiguous speech syllables whereas attention to the spectral (prosodic) properties of the same stimuli leads to activity in the right STS (Liebenthal et al., 2016). The present study demonstrates that pHVA is not only part of the WORD_ CORE network, but is also conjointly right activated and asymmetrical during the 3 Word-List tasks. These results suggest that pHVA would be involved in the tonal processing of word lists, underlying speech segmentation processing for each task (PROD, LISN and READ). Moreover, the rhythmicity of word lists processed by the right pHVA, would be the basis of the articulatory process which involves the left audio-motor loop, which is consistent with the recent neuroscientific literature supporting the use of musical training. Rhythmic stimulation related to the rhythm and intonation patterns of speech (prosody) has been revealed to improve auditory processing, prosodic and phonemic sensitivity in dyslexic children who perform poorly on tasks of rhythmic perception and perception of musical meters (Flaugnacco et al, 2015).

### The bilateral WORD_EXE network includes right visuospatial and attentional cortical areas supporting the executive aspects of the tasks

Brain areas constituting the second network were more related to the attentional processes conjointly shared by the three tasks, which is in accordance with the strong anticorrelation of the WORD-EXE network with the WORD-CORE network. In fact, in the three conditions, once the task was completed, subjects had to detect the transformation of a centrally displayed cross into a square and they were asked to press a button. Among these areas, SMA1, SMA2 and SMA3 overlapping the location of the supplementary frontal eye fields correspond, partially, to the dorsal frontal attentional network (Corbetta and Shulman, 2011). The fact that they were more activated on the right and more rightward asymmetrical during PROD, which was the most effortful task, is in line with a role in attentional control. The rightward asymmetry and activation of the precuneus regions, without any between-task difference suggest that it could be related to mental imagery triggered by the scramble version of the picture or to episodic memory encoding in reference to the list of the days and months (Cavanna and Trimble, 2006).

### General Conclusion and Perspectives

The present study, based on the fMRI analysis of 3 Word-List tasks performed by 144 healthy adult right-handers combined with the analysis of intrinsic resting-state connectivity in 138 of the same participants, makes itpossible to propose, for the first time, a model of the neural organization of Word-List processing during production, listening and reading tasks. This model posits that (1) action and perception circuits are interdependent and organised in networks, among which a trace of the learning modality is still present in the brain (2) the involvement of phonological action perception circuits such as the phonological working memory loop, in which articulatory gestures are the central motor units on which word perception, production and reading would develop and act according to the motor theory of speech (Liberman and Whalen, 2000) as revealed by the recruitment of leftward frontal and precentral areas together with temporo-parietal areas, and (3) the involvement of the right STS3 (pHVA), which is a prosodic integrative area, together with the left SMG1 which could reflect the intertwining between prosodic and phonemic information. The set of regions that constitutes WORD_CORE are available for download at http://www.gin.cnrs.fr/en/tools/.

### Ethics statement

The study has been approved by the Basse-Normandie rensearch ethics committee and has been performed in accordance with the ethical standards as laid down in the 1964 Declaration of Helsinki and its later amendments or comparable ethical standards.

### Informed consent

Informed written consent was obtained from all individual participants included in the study.

### Availability of Data And Material

The datasets analyzed in the current study to obtain the atlas are not publicly available because they were founded by the laboratory. They will be available when the laboratory completes all the publications. The Atlas is publicly available at http://www.gin.cnrs.fr/en/tools/.

## Authors’ Contributions

IH, LL and NTM wrote the article. LL, NTM and MJ completed the data analyses. NTM designed the paradigm and acquired the data and NTM and IH analysed the fMRI language runs, which are part of the BIL&GIN. All authors have contributed to the preparation and writing of the manuscript.

## Acknowledgements

The authors would like to thank Gael Jobard, Sandrine Crémona and Bernard Mazoyer for thoughtful comments and Gaelle Leroux for dedicated technical support.

